# Decoding visual colour from scalp electroencephalography measurements

**DOI:** 10.1101/2020.07.30.228437

**Authors:** Jasper E. Hajonides, Anna C. Nobre, Freek van Ede, Mark G. Stokes

## Abstract

Recent advances have made it possible to decode various aspects of visually presented stimuli from patterns of scalp EEG measurements. As of recently, such multivariate methods have been commonly used to decode visual-spatial features such as location, orientation, or spatial frequency. In the current study, we show that it is also possible to track visual colour processing by using Linear Discriminant Analysis on patterns of EEG activity. Building on other recent demonstrations, we show that colour decoding: (1) reflects sensory qualities (as opposed to, for example, verbal labelling) with a prominent contribution from posterior electrodes contralateral to the stimulus, (2) conforms to a parametric coding space, (3) is possible in multi-item displays, and (4) is comparable in magnitude to the decoding of visual stimulus orientation. Through subsampling our data, we also provide an estimate of the approximate number of trials and participants required for robust decoding. Finally, we show that while colour decoding can be sensitive to subtle differences in luminance, our colour decoding results are primarily driven by measured colour differences between stimuli. Colour decoding opens a relevant new dimension in which to track visual processing using scalp EEG measurements, while bypassing potential confounds associated with decoding approaches that focus on spatial features.

## 1. Introduction

Human scalp electroencephalography (EEG) and magnetoencephalography (MEG) are sensitive to synchronous activity in large neural populations, thus providing a macroscopic readout of brain activity. Whereas extracranial measures of brain activity carry the problem of source ambiguity (Koles 1998), it is nevertheless possible to compare scalp topography of brain activity to differentiate spatial patterns generated by subtly different configurations of neuronal populations (Stokes, Wolff, and Spaak 2015). Recently, studies have shown that the information across an array of electrodes can be used in multivariate analyses to decode neural information such as stimulus orientation, spatial location, spatial frequency, and motion direction (Bae and Luck 2019; Cichy, Ramirez, and Pantazis 2015; Foster et al. 2017; King and Dehaene 2014; Myers et al. 2015; e.g. Ramkumar et al. 2013; Sandhaeger et al. 2019; Stokes, Wolff, and Spaak 2015).

Fewer studies have successfully applied multivariate approaches to decode non-spatial features, such as colour. Decoding of visual-spatial features, which have typically been the focus in EEG decoding studies to date (such as orientation, location, and spatial frequency) is susceptible to contributions of spatial attention and/or eye movements, such as spatial biases in micro-saccades (van Ede, Chekroud, and Nobre 2019; Engbert and Kliegl 2003; Hafed and Clark 2002; Hollingworth, Matsukura, and Luck 2013; Mostert et al. 2018; Quax et al. 2019; Thielen et al. 2019). Therefore, the extension of decoding methods to features like colour, which are not defined by spatial parameters, provides an important validation of the approach by ensuring that decoding is based on the neural processing of feature-specific content. The demonstration of colour decoding from scalp EEG should therefore have important implications – affording novel ways to track perceptual representations in various contexts.

To date, a number of MEG (Hermann et al. 2020; Rosenthal et al. in press; Sandhaeger et al. 2019; Teichmann et al. 2019, 2020) and, to a lesser extent, EEG studies (Bocincova and Johnson 2018; Sandhaeger et al. 2019) have applied decoding approaches to distinguish among colour values upon visual presentation. We have built on these initial studies to further establish and characterise the properties of colour decoding from scalp EEG measurements. For example, it remains to be determined whether the relevant signals represent the sensory processing of colours and are independent from other associated factors, such as verbal labels or decision-making in tasks, or from subtle differences in luminance that can occur between rendered colours.

Like orientation, colour also follows an organisational structure in visual cortices (Engel, Zhang, and Wandell 1997; Kleinschmidt et al. 1996), and research has shown that it is possible to decode colour with functional MRI measurement with millimetre resolution (Brouwer and Heeger 2009) across a range of cortical visual areas (Brouwer and Heeger 2009; Persichetti et al. 2015). Therefore, finding that colour-related decoding follows parametric variations in stimuli (as opposed to binary or categorical boundaries) in a manner akin to decoding of stimulus orientation is an important step toward ensuring that decoding can reveal colour-specific neural information (in line with Rosenthal et al. in press). Further reassurance would come from the involvement of electrodes sensitive to visual processing to the decoding, at posterior sites posterior sites contralateral to the decoded stimulus (akin to recent MEG research; Hermann et al. 2020). Finally, the ability to decode the colours of multiple stimuli presented concurrently (as opposed to a single central stimulus in isolation, as in most prior colour decoding studies) would greatly add to the flexibility and analytical power of colour decoding as a tool in future studies.

In the present study, we therefore aimed to track parametric variations in colour processing in scalp EEG, from multi-item displays, while also qualitatively comparing ‘colour decoding’ to the more established decoding of stimulus orientation (in term of robustness, tuning profile, timing, spatial topography, and so on). To do so, we presented 30 participants with two sinusoidal Gabor gratings (one left, one right) that each contained a unique colour and orientation, and we required participants to attend to both features of both items (for a later memory test, which is beyond the scope of the current study). Using Linear Discriminant Analysis, we obtained reliable (and comparable) decoding of the colour and orientation of both stimuli. Our data also provide a clear case for a “visual” account of this decoding, by showing that colour decoding has a contralateral-posterior topography, and follows the parametric colour-coding space.

## 2. Materials & Methods

### 2.1 Participants

Ethical approval for this study was obtained from the Central University Research Ethics Committee of the University of Oxford. Thirty healthy adults (28.3 years old on average; ranging from 18 - 35 years old; 17 females) were included in this study after excluding two participants due to technical difficulties and incomplete data collection. All participants had normal or corrected-to-normal vision and reported no deficiencies in their colour vision. All participants were reimbursed £15 per hour for their time.

### 2.2 Visual display

Stimuli were generated using the Psychophysics Toolbox version 3.0.11 (Brainard 1997) in MATLAB 2014b (MathWorks, Natick, MA). The stimuli were presented on a 27-inch monitor (1920 × 1080, 144Hz; default gamma correction of 2.2). The monitor was integrated in a noise-proof booth with a glass barrier right in front of the display. Viewing distance was 90 cm from the monitor where participants were stabilised at a fixed position using a chin rest. Using a photodiode we established a 24 ms delay between the stimulus triggers received on the acquisition software and the visual presentation of the stimulus on screen. We corrected for this in our analyses accordingly.

### 2.3 Stimuli & design

The critical stimuli for this experiment were presented within the context of a visual working-memory task that ensured attentive viewing of the visual stimuli at encoding. Task details are presented here for clarity, but the analyses only focus on decoding the memory stimuli at encoding (for a full description of the task and participants’ behavioural performance, see: Hajonides et al. 2020). At the start of each trial, two Gabor stimuli with a diameter of 4.3 degrees of visual angle (one to the left, the other to the right of fixation, centred at ±4.3 degrees each) were presented simultaneously for 300 ms (**Fig. 1C**) on a grey background (defined RGB: 0.196 0.196 0.196; measured CIE L*a*b*: 15.18, 3.146, −3.822). The stimuli consisted of bilateral luminance-defined sinusoidal Gabor gratings and each had a unique colour and orientation. Colours were drawn from a CIELAB colour space, which is often used to define perceptually uniform stimulus sets. Forty-eight evenly-spaced colours were drawn from a circle in CIELAB colour space with a fixed lightness of L = 54 (**Fig. 1A**; centre is at a = 18, b = −8, radius =59).

**Fig. 1.**
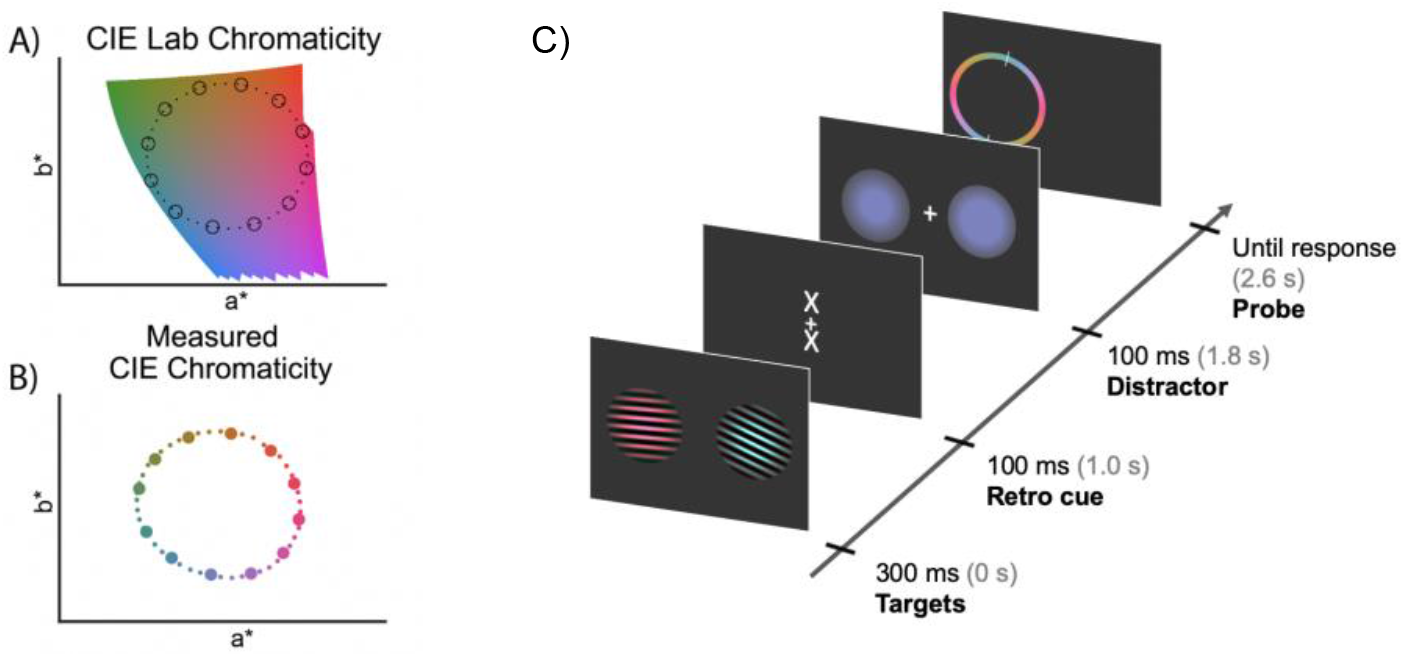
Stimulus characteristics and presentation. A) Forty-eight colours - depicted in small dots - were sampled from CIELAB colour space with a fixed lightness (“L”) of 54. During analysis, colours were binned into 12 evenly-spaced bins, depicted in circles. B) Measured CIE chromaticity a* and b* coordinates on the monitor using the Colorimeter. C) Participants were presented with two Gabor gratings for 300 ms with both colour and orientation information. Colour and orientation were varied independently, using circular spaces with 48 possible feature-values (indicated in panel A and B). Participants were required to reproduce the exact colour or orientation of either stimulus after a brief memory delay, this ensured attentive encoding of the stimuli. The memory delay contained two events: a retro cue, and a visual distractor that consisted of a bilateral colour circle or two identical oriented Gabor gratings. We focus our analyses on the 1000 ms after onset of the gratings. The schematic shows an example trial prompting a report for the colour of the left grating. Stimulus durations are shown next to the tick marks along with in brackets the onset time after the start of the trial. See Hajonides et al. (2020) for the full details of the task design.

To get a precise measurement of the obtained luminance of our stimuli on our monitor, forty-eight rendered colours were measured post-hoc using a CRS ColorCal KMII Colorimeter (Cambridge Research Systems, Kent, UK). Luminance and CIE chromaticity (in CIE xyY coordinates) were measured on a uniform colour patch containing only the average colour of the sinusoidal grating as an approximation of the grating colour with varying intensity. CIE xyY coordinates were converted to CIELAB coordinates using a d65 white point (Fig. 1B). Deviations of the rendered colours from the CIELAB colours can be caused through various ways. For example, CIELAB colours were converted using a linear colour conversion algorithm (Image Processing Toolbox, MATLAB, The MathWorks, Inc., Natick, MA) with an industry standard d65 white point. Without monitor calibration, the sampled RGB colours could deviate from originally defined CIELAB colours. Also, not all colours can be rendered by computer monitors. This is likely to impose small non-systematic imprecisions on the generation of individual colours. Finally, as previously described, there was a glass barrier between the monitor and the participant. The glass barrier could be introducing reflections, resulting in light loss. We added these deviations to our analysis to assess the relative contributions of stimulus rendering offsets and to map out (and rule out) potential alternative explanations for our colour decoding (such as decoding of luminance differences, rather than colour differences, between our stimuli).

In total, 48 different colours and 48 different orientations were presented, with feature values ranging from 3.75 to 180 degrees in evenly spaced steps. Orientation and colour values were initially allocated randomly within each block. However, to minimise trials in which the two items had the same or very similar colour or orientation values, we redrew the colour and orientation values of all trials in a block when more than 2 trials were allocated two items with colour or orientation values within the same bin (out of 12 bins). To increase the amount of samples for feature, we binned the 48 colour and orientation features into 12 - equally spaced – bins (**Fig. 1AB**). To match the orientation space more closely, we opted for a symmetrical colour probe that included all colours in the 0 – π range of the probe.

At the time of stimulus presentation, participants were instructed to attend to and remember both the colour and the orientation of both gratings. At the end of each trial, participants were probed to report a single feature from one of the two items. Which feature and item would be probed was unpredictable at encoding. During the 2400-ms memory delay, two events occurred: (1) 1000 ms after encoding onset a cue could provide information about the item or feature dimension that would be probed and (2) 1800 ms after encoding onset, a bilateral colour or orientation distractor was presented to distort visual processing. These trial details are not critical to the current investigation, which focuses almost exclusively on the encoding phase of the experiment. More detailed description of the experiment from which we here re-analysed the EEG data can be found in Hajonides et al. (2020).

### 2.4 EEG recording

The EEG signal was acquired using a 10-20 system using 61 Ag/AgCl sintered electrodes (EasyCap, Herrsching, Germany). The data were recorded using a Synamps amplifier and Curry 8 acquisition software (Compumedics NeuroScan, Charlotte, NC). An electrode placed behind the right mastoid was used as the active reference during acquisition. Offline, data were re-referenced to the average of the left and right mastoids. The ground was placed above the left elbow. Bipolar electrooculography (EOG) was recorded from electrodes above and below the left eye and lateral of both eyes. EEG data were digitised at 1000 Hz with an anti-aliasing filter with a cut-off frequency of 400 Hz. Impedances were kept below 7kΩ.

### 2.5 EEG pre-processing

EEG data were analysed in MATLAB 2017a using FieldTrip (Oostenveld et al. 2011) in conjunction with the OHBA Software Library (OSL; https://ohba-analysis.github.io/). After importing, the data were epoched from −300 ms to 1000 ms around presentation the presentation of the Gabor gratings (ft_redefinetrial) and re-referenced to the average of both mastoids (ft_preprocessing). Raw epoched data were downsampled to 100 Hz, and stored for subsequent analyses (analysis scripts and data is made available online at OSF https://osf.io/j289e/?view_only=b13407009b4245f7950960c34a5474a6).

Subsequently, we used the EOG to identify trials in which participants blinked and might have missed the visually presented stimuli. The vertical EOG was baselined between 300 ms to 100 ms before stimulus presentation. Trials that exceeded half of the maximum voltage elicited by blinks (half of 400 μV) within the timeframe of 100 ms prior to and 100 ms post stimulus presentation were marked and later discarded. Additional eye-related artefacts were identified using Independent Component Analysis (ICA; ft_componentalanalysis) by applying the FastICA algorithm (Hyvärinen 1999) to the full EEG layout. Components were rejected if they had a correlation of *r* > 0.40 with the EOG electrodes (on average 2.7 components per participant, and a maximum of 4 components). In addition to trials where participants blinked at the moment of stimulus presentation, trials with high within-trial variance of the EEG broadband signal were removed using a generalised ESD test at a 0.05 significance threshold (Rosner 1983; implemented in OSL), removing 2.48 % ± 2.18 % (mean ± standard deviation) of all trials.

### 2.6 Classification on EEG data

To focus our decoding on visual processing, we restricted our main analyses on the 17 most posterior electrodes (P7, P5, P3, P1, Pz, P2, P4, P6, P8, PO7, PO3, POz, PO4, PO8, O1, Oz and O2; as in Michael J Wolff et al. 2017).

To make class predictions we employed Linear Discriminant Analysis (LDA) using the Scikit-learn toolbox (Pedregosa et al. 2011) and estimated the likelihood for all classes. LDA is a multi-class decoding algorithm that performs well for neuroscientific time-course data (for a comparison of classifiers see Grootswagers, Wardle, and Carlson 2017).

To increase the sensitivity of our decoding analysis, we capitalised on both spatial and temporal patterns for decoding (Grootswagers, Wardle, and Carlson 2017; Michael J. Wolff et al. 2020). To do so, we used a sliding-window approach and concatenated topographical patterns from t_0_ up to t_−19_ (20 steps of 10 ms, ranging over 200 ms) into a single vector. This increased the amount of features in the LDA classifier from 17 (17 posterior electrodes) to 340 (17 electrodes × 20 time steps). To avoid baselining issues and to utilise dynamic activity rather than stable brain states, we subtracted the mean activity across the 20 time points within each time window for each channel and trial separately. This analysis method thus exploits embedded informative temporal variability, reduces noise by combining data over time, and circumvents baselining issues (Grootswagers, Wardle, and Carlson 2017; Michael J. Wolff et al. 2020). In a supplementary analysis, we confirmed that similar (be it less robust) colour and orientation decoding could be obtained when decoding based on purely spatial patterns, on a timepoint-by-timepoint basis.

Principal component analysis was applied to the data to capture 95% of the variance in the data (sklearn.decomposition.PCA). Trial feature labels were binned into 12 classes (each subtending 12/π for orientation and 6/π for colour; numpy.digitize). These data were split into train-and-test sets using 10-fold stratified cross-validation (sklearn.model_selection.RepeatedStratifiedKFold). Next, training data were standardised (sklearn.preprocessing.StandardScaler) and after the same standardisation was applied to the test data we fit the LDA to the training data and labels (sklearn.discriminant_analysis.LinearDisciminantAnalysis; using singular value decomposition and a default threshold for rank estimation of 0.001). Likelihoods for each class were estimated after training and testing the LDA classifier separately for each time point, features (colour and orientation from left and right sides), and participant. The resulting likelihoods were centred so that the predicted class was always in centre surrounded by neighbouring feature classes. These predicted-class likelihoods were convolved with a cosine (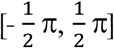 for orientation and [− π, π] for colour) to provide a single measure of evidence. This serves to convert two-dimensional tuning curves and obtain a single measure of evidence of each time point. For additional analyses, where we looked at dissimilarity matrices, no cosine convolution was applied. Training and testing were performed separately for left and right features and only at the final stage we averaged across both sides.

To confirm the posterior locus of decodability, we also ran a searchlight decoding analysis (Kriegeskorte, Goebel, and Bandettini 2006), iteratively considering a small group of electrodes. To this end, we applied the same temporal classification on the voltages between 150 and 350 ms, but this time only considered the data from a given electrode together with the data from its immediately adjacent (left/right/above/below) neighbours.

Building on RSA (representational similarity analysis; Kriegeskorte, Mur, and Bandettini 2008) we additionally investigated how our measured differences in luminance (L*) and colour (a*b*) each contributed to the identified parametric decoding of colour. RSA allowed us to examine relative contributions of different neural codes. The preceding analyses assumed a perfectly circular organisation of colours (**Fig. 1A**) with identical tuning curves for each colour. In practice, rendered colours slightly violate this hypothesised organisation. To test the relative contribution of luminance and colour in our data, we therefore constructed ‘dissimilarity models’ for both features using the measured Colorimeter data. For measured luminance (L*), a one-dimensional variable, we characterised the difference in luminance of each of the 12 colour bins as the absolute difference of one colour bin with all other colour bins. Doing so for each colour, we obtained a 12-by-12 matrix containing differences of measured luminance of each colour with all other colours with a value of zero across the diagonal. Similarly, we generated a model characterising differences in measured a*b*. Here, we used Euclidean distance between measured a*- and b*-coordinates of each colour bin and all other colour bins as a proxy of colour dissimilarity. A third model acted to compare these two models to the circular model (that used the desired CIELAB colour space as its basis) described above. All models were z-scored. Subsequently, all three models were multiplied - and subsequently summed - with LDA evidence of every colour bin for each of the 12 possible classifier predictions for every subject, using the spatial-temporal decoding method described above. Higher scores indicate higher similarity between the patterns in the data and those in the model.

### 2.7 Statistical analysis

For the analysis of the classifier predictions, cosine-convolved evidence for each decoded feature-dimension (colour, orientation) was compared to data with labels that were randomly shuffled prior to training the classifier for each time point using a cluster permutation test (mne.stats.permutation_cluster_test; Gramfort et al. 2014) with 10000 iterations and a F-statistic threshold of 2.045 (p < .05 with 29 degrees of freedom). Cluster-based permutation tests were also applied to the similarity measure obtained in the RSA analysis. Here, we compared similarity of the measured a*b* model and that of the defined CIELAB model against the luminance model to ask whether we could decode CIE chromaticity over and above measured differences in luminance. For completeness, we also tested whether measured luminance could be significantly decoded, for this we tested similarity scores of the luminance model against similarity scores of a 12-by-12 matrix of z-scored random values multiplied with the data.

For further characterisation of the data, we also ran several analyses with data over the 150-350 post-stimulus time-window in which both colour and orientation decoding were particularly prominent. We first used this window to assess the reliability of the LDA likelihoods for each of the 12 colours and orientations separately. For each bin we tested decoding against zero using a one-tailed one-sample t-tests, and report both the uncorrected results and the results after applying a Bonferroni correction for multiple comparisons. We also used the data from the same window to investigate the parametric coding spaces for colour and orientation by applying multi-dimensional scaling (MDS; sklearn.manifold.MDS; Kruskal, 1964) to a matrix with rows for decodable colour or orientation and columns for classifier predictions. This matrix was inverted so that the diagonal had the smallest likelihoods. We ran MDS with 1000 initialisations and a maximum of 1000 iterations to find the configuration with the lowest stress score for a two-dimensional embedding of the 12 colours and orientations. Finally, we varied the number of participants and trials/participant we included in our analysis. Doing so, we randomly sampled a subset of participants (5-30) and the first *n* trials/participants (where *n* ranged from 96 to 912 in steps of 48), we took this approach to simulate having run a shorter experiment. Subsequently, we evaluated the cosine-convolved classifier evidence resulting from spatial-temporal decoding between 150 ms and 350 ms for colours and orientations, averaged over left and right stimulus presentations. To ensure random sampling we drew each subset of participants over 100 permutations and averaged over the results. In contrast to other analyses, a 5-fold cross validation was used to deal with the full range of trial numbers. The resulting matrix was slightly smoothed using a gaussian kernel with a sigma of 0.5 (scipy.ndimage.gaussian_filter), where edge artefacts were avoided by replicating the nearest data point.

## 3. Results

### 3.1 Colour can be decoded from scalp EEG, and is comparable to orientation decoding

We extracted EEG data from the 17 posterior electrodes ranging from 300 ms prior to the presentation of the two gratings to 1000 ms after stimulus presentation onset. Average ERP traces revealed similar voltage profiles across the range of tested colours (**Fig. 2A**, left) and orientations (**Fig. 2A**, right). An LDA classifier was able to uncover a clear reconstruction of the presented colours and as well orientations (**Fig. 2B**; averaged over left and right stimulus presentations). To increase sensitivity and to ease visualisation, we reduced these tuning-profiles from **Fig. 2B,** to a single decodability value per time point, by convolving the tuning profiles with a cosine function (van Ede et al. 2018; M. J. Wolff et al. 2019; as in Michael J Wolff et al. 2017). Decoding time-courses are depicted in **Fig. 2C**. We found highly significant decodability for both visual features. Colour decoding first reached significance after 115 ms and remained significant until 875 ms. Orientation decoding first reached significance after 135 ms and remained significant until 945 ms. Colour decoding could also be demonstrated for the processing of the visual distractors, which were not attended and did not require a behavioural report (**Fig. S1**)

**Fig. 2.**
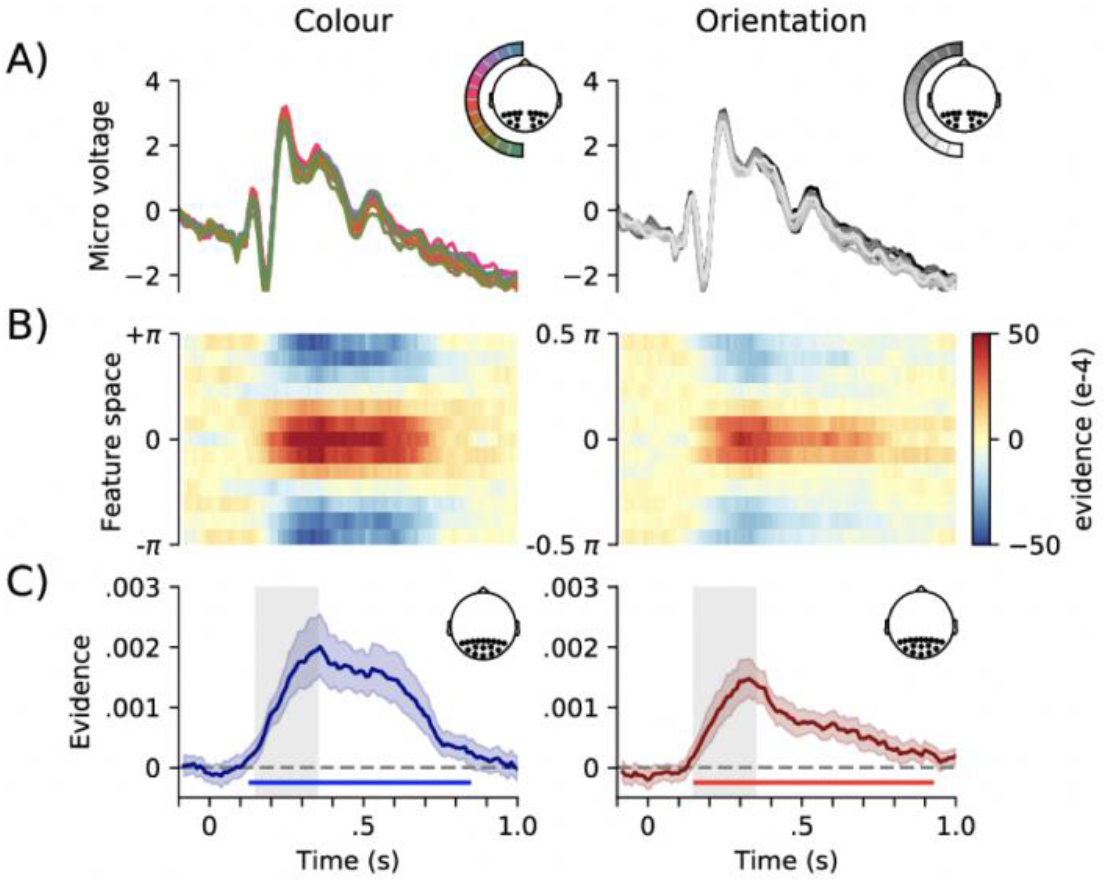
ERPs and decoding performance for colours and orientations. A) Event-related potentials for the 7 posterior electrodes contralateral to the presented feature (see inset), averaged over left and right, locked to stimulus presentation. Displaying micro voltage for all twelve different orientations or colours. B) Two-dimensional, decoding tuning curves with the likelihood for target colour or orientation (value “0”) and all neighbouring values of theta. For visualisation purposes, a 13^th^ bin was added at the bottom on the diagram mirroring the top bin. These multivariate analyses were done using only the 17 posterior electrodes (see inset 2C). C) Mean cosine-convolved evidence with red lines for colour, blue for orientation. Cluster-permutation corrected significant time points are indicated with horizontal lines. Grey-shaded area between 150 to 350 ms is used for subsequent analyses. These multivariate analyses were done using only the 17 posterior electrodes (see inset 2C). Error bars show 95% confidence intervals, calculated across participants (n = 30).

### 3.2 Colour decoding is driven by visual signals with a contralateral-posterior topography

To understand what contributed to our ability to decode colour, we next ran a searchlight decoding analysis (Kriegeskorte, Goebel, and Bandettini 2006) across all sensors (van Ede et al. 2019; Michael J. Wolff et al. 2020). This demonstrated that classifier evidence (in the 150 to 350 ms post-stimulus period in which we found robust decoding; grey-shaded windows in **Fig. 2C**) peaked in posterior electrodes, with primary contributions from electrodes contralateral to the decoded stimuli. This was the case both for decoded colour (**Fig. 3A**), and orientation (**Fig. 3B**). This provides compelling evidence for the “visual” nature of this decoding (as opposed to, for example, decoding distinct “verbal labels” associated with distinct colours).

**Fig. 3.**
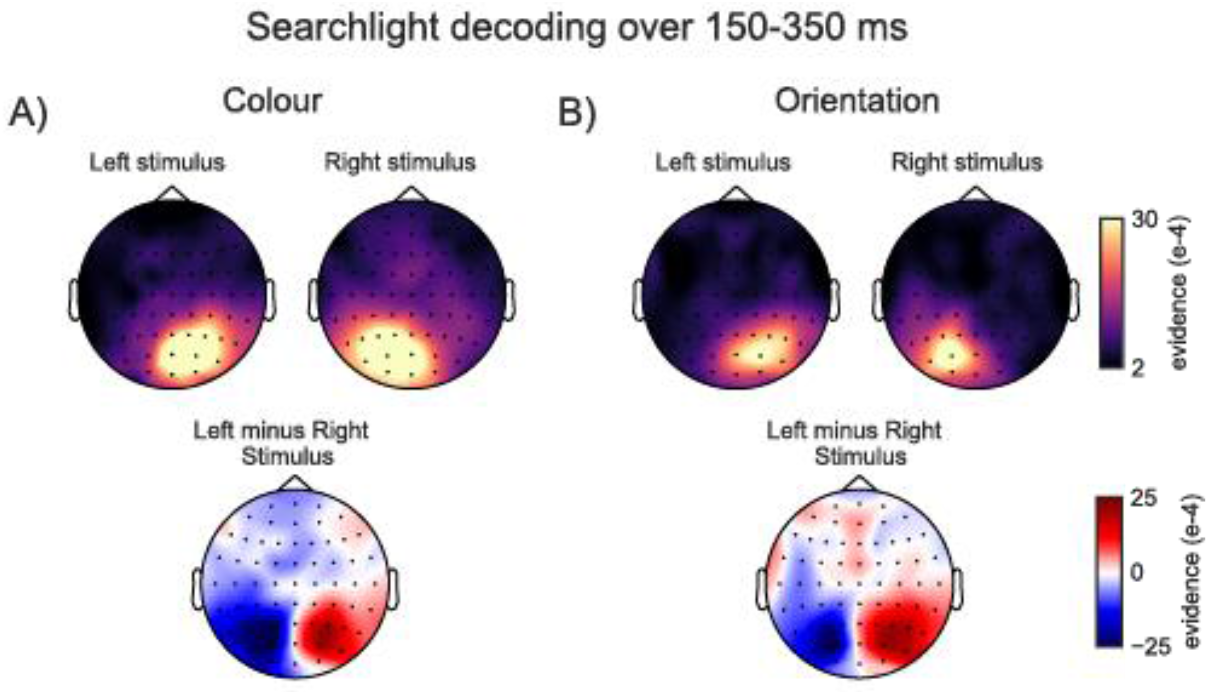
Colour and orientation decoding are primarily driven by posterior electrodes contralateral to the decoded stimulus. Top row topographies show which electrodes show highest mean cosine-convolved evidence for A) colours and B) orientations, of stimuli that were presented on the left or on the right of fixation. Posterior contralateral electrodes show highest evidence. Bottom panels show the difference between left-stimulus and right-stimulus decoding topographies, highlighting the lateralisation of the evidence, and the comparable lateralisation obtained for colour and orientation decoding.

### 3.3 Colour decoding is parametrically organised

To test how well each of the feature-value bins could be decoded (relative to all other feature-value bins within the colour and orientation spaces), we analysed mean cosine-convolved LDA likelihoods in the data pooled between 150-ms and 350-ms window that showed robust decoding of both features (indicated in grey in **Fig. 2C**). Decoding, as a function of the decoded feature-values, is shown in **Fig. 4A**. For colour bins, all 12/12 colours could be decoded significantly against a *p* < .05 threshold (mean t_29_ = 4.523; min = 2.440; max = 6.175), and for orientation 11/12 orientations could be decoded significantly against an uncorrected single-tailed threshold (mean t_29_ = 4.096; min = 1.585; max = 7.390). After applying Bonferroni correction, 9/12 colour bins were significant and 8/12 orientations. Using multi-dimensional scaling we revealed the representational space in two dimensions, showing a circular, or oval, configuration for both colour and orientation bins (**Fig. 4B**). For orientation, the cardinal orientations (0, 90, and 180 degrees) showed higher decoding than the off-cardinal decoding. Interestingly, due to some non-linearities in the circular space, we observe some colours (pink, red, orange) in one part of the representational space and other colours (purple, blue, green) on the other (see also Hermann et al. 2020).

**Fig. 4.**
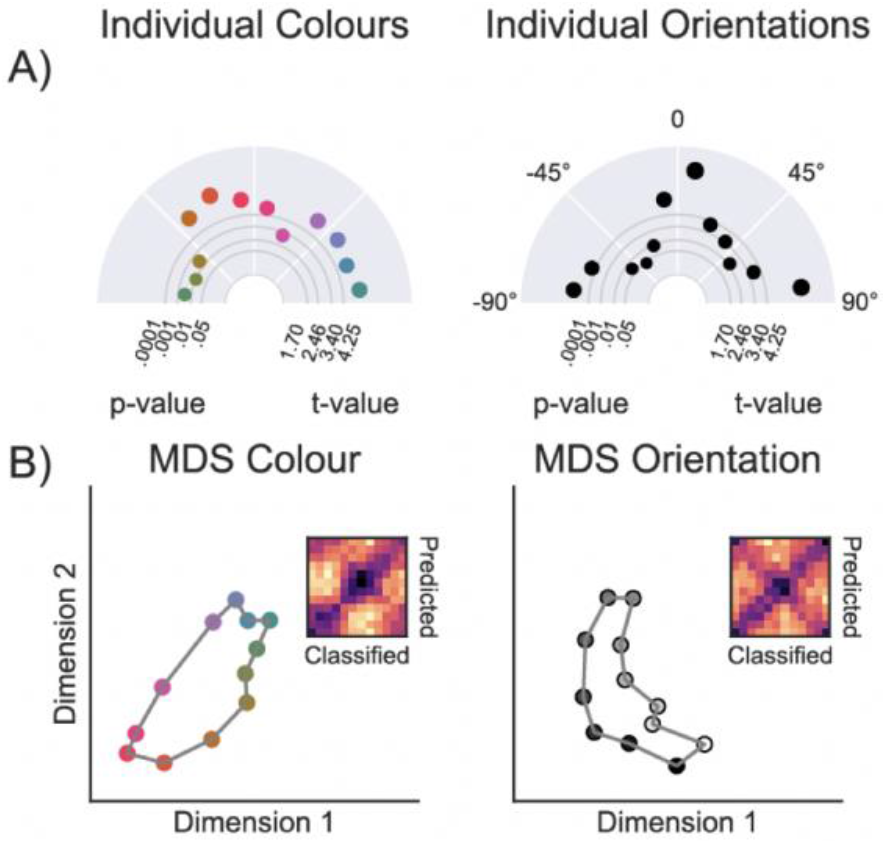
Decoding of individual features between 150 and 350 ms. A) This polar plot illustrates the decoding evidence of 12 individual colours and orientations. The distance from the centre depicts the t-value of testing the cosine-convolved tuning curve for a single feature, left and right combined, across participants, relative to zero. The size of the dots mirrors the magnitude of the evidence for that feature B) Two-dimensional visualisation of the similarity matrix in which we observed a circular configuration for colour features. Dissimilarity matrices show feature (colour or orientation) on the y-axis and distance between bins of the tuning curve on the x-axis.

### 3.4 Decoding is sensitive to luminance (L*) differences but primarily driven by difference in colour (a*b*)

Colours generated for this experiment were drawn from a plane in CIELAB with identical (desired) lightness, and with equal distances between the colours in the circular colour landscape (Fig. 1A). Nevertheless, because of slight luminance and colour differences generated during the conversion to RGB colours and rendering process on the monitor, measured colour properties were slightly different. In the next step, we therefore used the physically observed measured luminance (L)* and measured colour (a*b*) to dissect their relative contributions to the identified colour decoding.

To this end, we created three “models” (**Fig. 5A**) and used RSA to ask how well each of them independently fit the empirical EEG data. We observed that measured differences in luminance (L*; Luminance model in **Fig. 5A**) between the 12 colour-bins could be decoded from the EEG signal yielding a significant cluster between 115 - 765 ms (**Fig. 5B**). Critically, however, in addition to the luminance model, we also compared the data to the dissimilarity matrix of the measured colour values (a*b*; **Fig. 5A**), as well as the (highly similar) CIELAB model used in the previous analyses. This revealed that the colour model and the CIELAB model each fit the observed data significantly better than the luminance model, with significant clusters between 155 - 855 ms and 165 - 735 ms, respectively (indicated by the increased line width in the horizontal cluster-significance lines in **Fig. 5B**). This is also appreciated by comparing the three models to the empirical dissimilarity matrices, at several time slices, in the bottom of Fig. 5B. These data confirm that the empirical dissimilarity matrix in the EEG data is more similar to the colour and CIELAB models, than to the model constructed from measured luminance values, and that this is so consistently across time. Thus, although unintended luminance differences between our stimuli may have made some contribution, the vast share of our colour decoding appears to be driven by colour(a*b*) differences between our stimuli.

**Fig 5.**
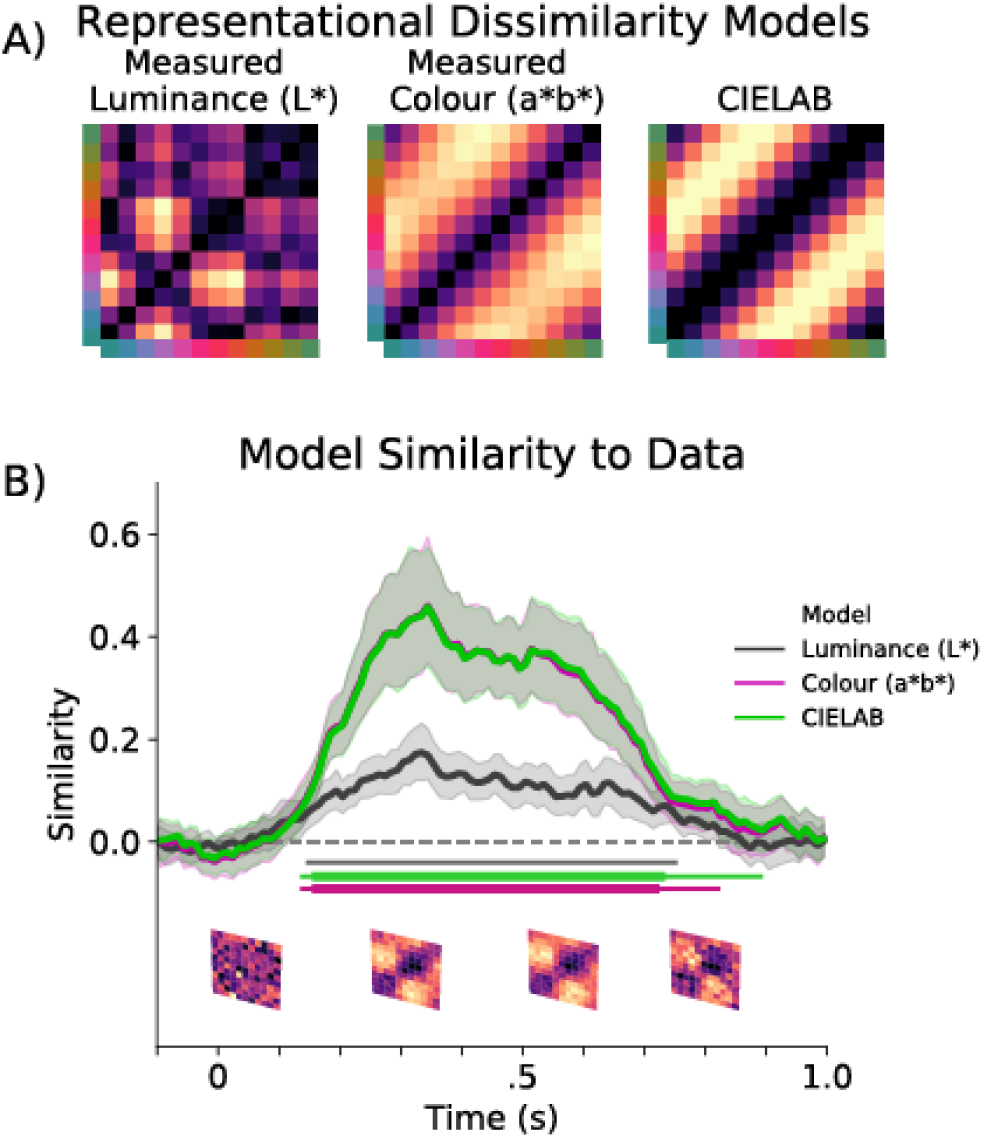
Decoding measured luminance and a*b* differences between stimuli. A) Z-scored dissimilarity models representing the measured differences is luminance (L*), Euclidean distances between 12 pairs of measured a*b* values, and a CIELAB model that assumes equal dissimilarity between all neighbouring colours (and that was used in all other analyses). B) Similarity between dissimilarity models and the LDA evidence in the EEG data for each time point between all 12 colour bins. Going from left to right, the matrices below the x-axis show the average LDA evidence between −100 to 150 ms, 150 to 400 ms, 400 to 650 ms, and 650 to 900 ms post stimulus onset. Clusters of significant differences between a random model and measured a*b* (purple), measured luminance model (black) and CIELAB (green) are indicated with the coloured horizontal lines below the similarity time courses. Larger line width of the horizontal green and purple lines indicate cluster-corrected differences with luminance evidence. Error bars show 95% confidence intervals, calculated across participants (n = 30).

### 3.5 Dependence of decoding on trial and participant numbers

The current study included 30 participants who each completed over 900 trials. To provide guidance on “how many” participants and trials would typically be required to achieve robust decoding, we ran the following analysis. We randomly sampled *x* participants (in 100 permutations) and, for each included participant, we incrementally increased the number of trials used for our decoding analysis, always including the first *y* trials of the session (as if we had only collected data up to this trial). For each combination of # participant by # trials per participant, we assessed feature decoding between 150 to 350 ms after stimulus onset, and plotted the resulting matrix (**Fig. 6**). Both the number of trials and the number of participants have a strong effect on the reliability of feature decoding. It is evident that both sources of statistical power influence our decoding scores.

**Fig. 6.**
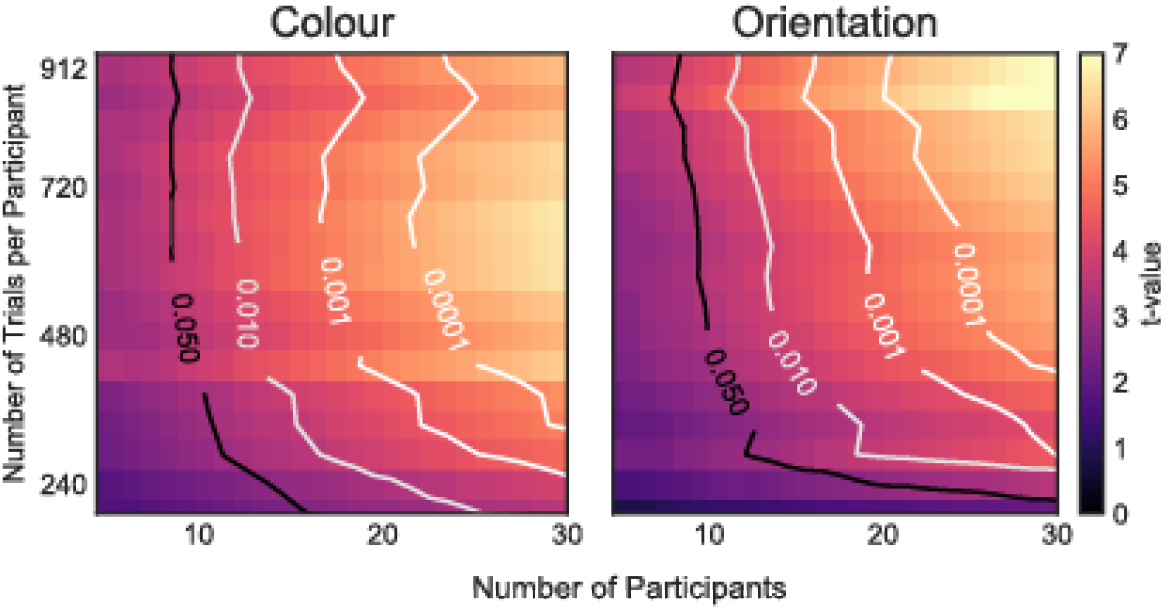
Feature decoding as a function of trial and participant numbers. Average cosine-convolved evidence of data between 150 and 350 ms for the average of left and right feature decoding. Contours mark the .05, .01, .001, and .0001 p-value thresholds for n-1 degrees of freedom.

## 4. Discussion

We investigated the ability to decode visual colours from scalp EEG measurements. We built on other recent studies that have employed colour decoding in scalp EEG (Bocincova and Johnson 2018; Sandhaeger et al. 2019) and MEG (Hermann et al. 2020; Rosenthal et al. in press.; Sandhaeger et al. 2019; Teichmann et al. 2019, 2020) and extended these in several ways. We show that we can reliably classify four simultaneously presented visual features – two colours and two orientations – from posterior EEG recordings using multi-class LDA. Follow-up analyses revealed that activity in posterior electrodes contralateral to the decoded stimulus were the primary contributors to the decoding of both features, suggesting that visual sensory processing was the main source of decodable signals, ruling out alternative explanations of colour decoding, such as verbal labelling. Furthermore, colour decoding followed a colour circle in representational space, speaking to the parametric nature of visual colour decoding. Finally, by having both orientation and colour information present in our stimuli, we were able to decode both. This revealed how the decoding of colour is comparable in magnitude, and if anything even clearer, to the decoding of orientation in a visual stimulus.

The current results are in line with, and nicely build on, several other recent non-invasive human electrophysiological studies showing an increase in classifier evidence following visual presentation of coloured stimuli (Hermann et al. 2020; Rosenthal et al. in press; Sandhaeger et al. 2019; Teichmann et al. 2019, 2020). One critical difference between the current work and these previously described studies is that we here used a 61-electrode EEG setup rather than a MEG system which typically contains a much larger number of sensors. Though MEG and EEG are sensitive to similar types of activity (but see Lopes da Silva 2013; Malmivuo 2012), demonstrating that it is also feasible to decode colour in scalp EEG is an important advance. This is because EEG has lower spatial resolution than MEG and because EEG is more accessible as a technique to labs around the world. Adding to this particular advance, our data further reveal that it is possible to decode parametric colour values in multi-item displays with lateralised stimuli - which allowed us to demonstrate a predominant contralateral topography - and that decoding of colour is not necessarily inferior (at least in our data) to the decoding of stimulus orientation.

To increase sensitivity, in the current work we capitalised on both spatial and temporal activity patterns for classification (see Grootswagers, Wardle, and Carlson 2017; Michael J. Wolff et al. 2020). However, to ensure that our results did not critically depend on this choice, we also ran all of our analysis using a more basic decoding approach whereby we relied only on spatial patterns for decoding; running the decoding analysis separately for each time point after stimulus onset. While this analysis was, as expected, less sensitive, we could nevertheless demonstrate that it was still possible to significantly decode visual colour from purely spatial patterns in the EEG data; that colour and orientation decoding remained highly comparable in magnitude; and that colour and orientation decoding each had a contralateral-posterior topography and conformed to a parametric coding space (**Fig. S2-5**).

Perceived colour and luminance depend on many factors, including monitor setup, local stimulus contrasts, and individual biology (Webster et al. 2000). While it is therefore difficult to disentangle fully the respective contributions of each of these factors to our ability to decode colour, our primary aim was to demonstrate its primary source and usefulness in cognitive neuroscientific research. Furthermore, to characterise the relative contributions of subtle difference in luminance from the desired differences in colour empirically, we performed post-hoc measurements of each and used RSA to estimate their relative contribution to our neural colour decoding results. Though we observed significant evidence of luminance differences between the rendered colour stimuli, we also found significantly stronger evidence for the decodability of both the intended (CIELAB) and the physically measured colour differences between rendered stimuli. We further observed a clear parametric coding space that conformed to the circular colour space (that was defined for colour, not for luminance). Furthermore, recent MEG research described how luminance and hue may modulate brain activity independently (Hermann et al. 2020), providing further validation that is it possible to decode colour, independently from luminance.

The ability to decode colour, as a reliable proxy of detailed visual processing, is important not only because it opens a new dimension through which to track detailed visual processing on the basis of scalp EEG measurements. It may also prove instrumental to by-pass a fundamental limitation faced by other decodable features such as spatial location, orientation, or spatial frequency. Colour, being a non-spatial parametric feature, should be much less susceptible to confounds associated with visual-spatial processing, such micro-saccades (van Ede, Chekroud, and Nobre 2019; Engbert and Kliegl 2003; Hafed and Clark 2002; Hollingworth, Matsukura, and Luck 2013; Mostert et al. 2018; Quax et al. 2019; Thielen et al. 2019).

## 5. Conclusion

In conclusion, like several other recent studies, we show that colour decoding is possible from scalp EEG measurements. Building on this related recent work, we have shown that this colour decoding reflects visual processing with a clear posterior-contralateral topography; that it conforms to a parametric colour-coding space; that it is possible in multi-item display; and that it is comparable to the decoding of stimulus orientation. This opens a relevant new dimension in which to track visual processing using scalp EEG measurements, while bypassing potential confounds associated with conventional decoding approaches that focus on spatial features.

## CRediT authorship contribution statement

**Jasper E. Hajonides:** Conceptualization, Methodology, Investigation, Formal analysis, Writing - Original Draft, Writing - Review & Editing, Visualization, Funding acquisition; **Anna C. Nobre:** Supervision, Conceptualization, Methodology, Writing - Original Draft, Writing - Review & Editing, Funding acquisition; **Freek van Ede:** Supervision, Conceptualization, Formal analysis, Writing - Original Draft, Writing - Review & Editing; **Mark G. Stokes:** Supervision, Conceptualization, Methodology, Investigation, Writing - Original Draft.

## Acknowledgements

We would like to thank Sammi Chekroud for his help with the experimental work as well as Hannah Smithson and Allie Hexley for lending us their colorimeter equipment.

This research was funded by an ESRC Grand Union studentship and the Scatcherd European Scholarship awarded to J.E.H., a Marie Skłodowska-Curie Fellowship from the European Commission (ACCESS2WM) and an ERC Starting Grant from the European Research Council (MEMTICIPATION, 850636) to F.v.E., and was supported by a James S. McDonnell Foundation Scholar Award (220020405), an ESRC grant (ES/S015477/1), the Medical Research Council Career Development Award (MR/J009024/1), and a Biotechnology and Biological Sciences Research Council award (BB/M010732/1) to M.G.S., as well as a James S. McDonnell Foundation Understanding Human Cognition Collaborative Award (number 220020448), and a Wellcome Trust Senior Investigator Award (104571/Z/14/Z) to A.C.N., and was supported by the NIHR Oxford Health Biomedical Research Centre. The Wellcome Centre for Integrative Neuroimaging is supported by core funding from the Wellcome Trust (203139/Z/16/Z). The funders had no role in study design, data collection and analysis, decision to publish or preparation of the manuscript.

## Supplementary materials

During the delay period, participants were presented with a bilateral distractor for 100 ms which consisted of two coloured discs (of the same colour) on half of the trials or two oriented Gabor gratings (of the same orientation) on the other half. Sixteen evenly spaced colour and orientation features were defined for the distractor, which were binned into 8 bins for the following decoding analyses to ensure a similar number of trials per feature. We epoched the data from 400 ms prior to distractor onset to 800 ms post distractor onset. The same spatial-temporal decoding methods described in the manuscript were applied to the neural data. Classifier evidence for colours was an order of magnitude higher than that observed for items in the encoding frame. This may be due to the fact that the distractors only contained a single feature that was identical for the left and right side.

**Fig. S1.**
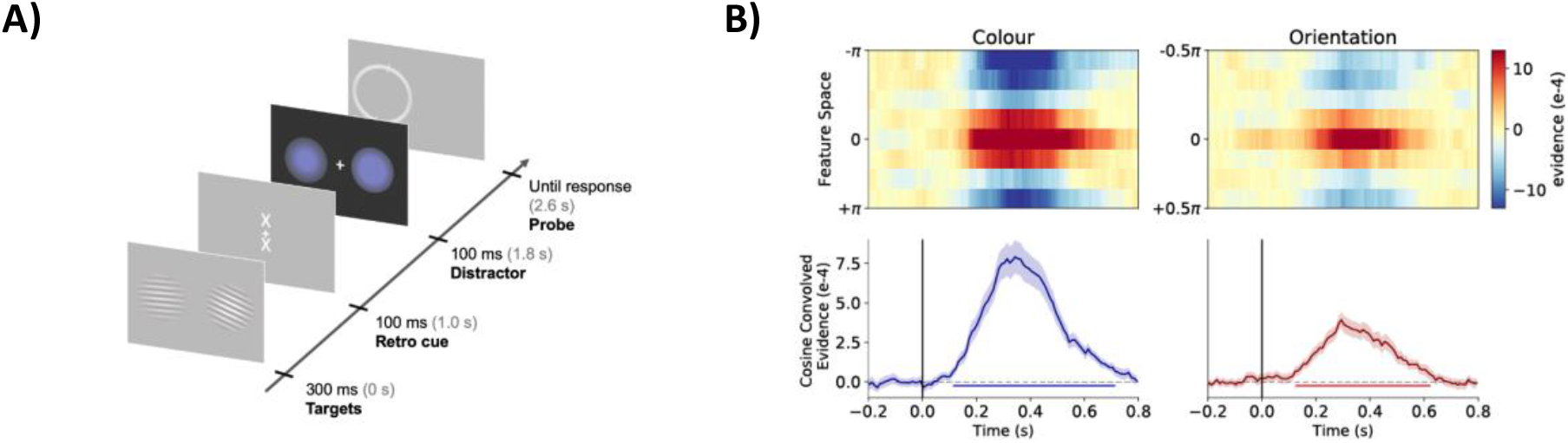
Decoding the colour and orientation of the visual distractors. A) On every trial, participants were presented with a bilateral array of distractors that consisted of a single feature. On a randomly selected half of trials this was a colour and on the other half of trials this was an orientation. The decoding analyses for the distractor epoch were carried out in the same way as in the encoding epoch, colour and orientation features were binned into 8 stimulus bins and decoding analyses were applied in an identical fashion for the distractor epoch. B) Mean cosine-convolved evidence with red lines for colour, blue for orientation. Two-dimensional, decoding tuning curves with the likelihood for target colour or orientation (value “0”) and all neighbouring values of theta. Error bars show 95% confidence intervals, calculated across participants (n = 30).

When applying LDA to the data from the 17 posterior electrodes for each time point, we do not reduce the dimensionality using PCA. Other preprocessing steps are identical. Single-time point LDA decoding shows that colour decoding was first significant after 125 ms, until 595 ms (**Fig. S2**). Orientation decoding was only significant after 185 ms, until 625 ms.

**Fig. S2.**
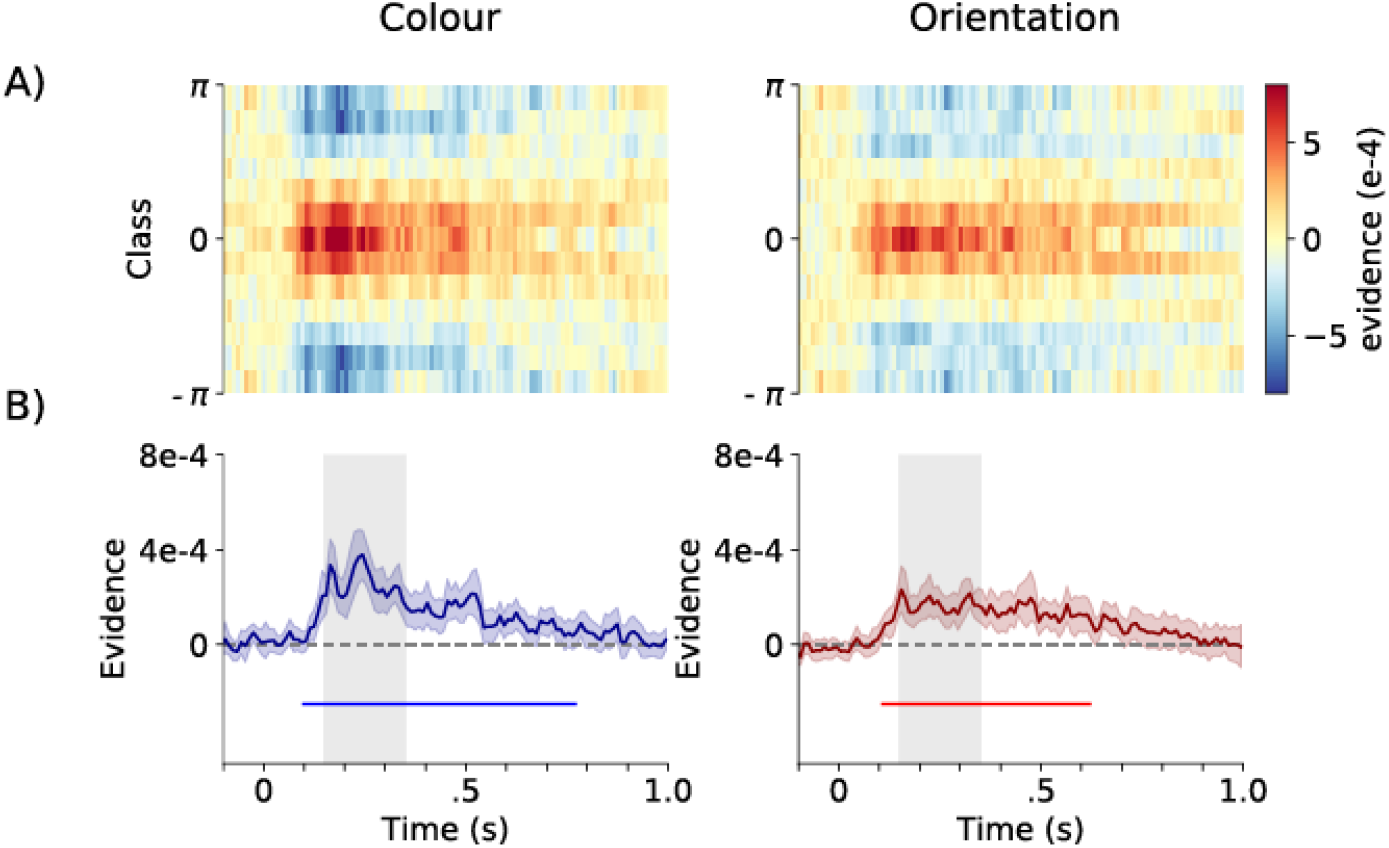
Decoding performance for colours and orientations with single time-point decoding. A) Two dimensional tuning curves with the likelihood for target colour or orientation and all neighbouring values of theta. B) Mean cosine-convolved evidence with red lines for colour, blue for orientation. Cluster-permutation corrected significant time points are indicated with horizontal lines. Grey-shaded area between 150 to 350 ms is used for subsequent analyses. Error bars show 95% confidence intervals.

**Fig. S3.**
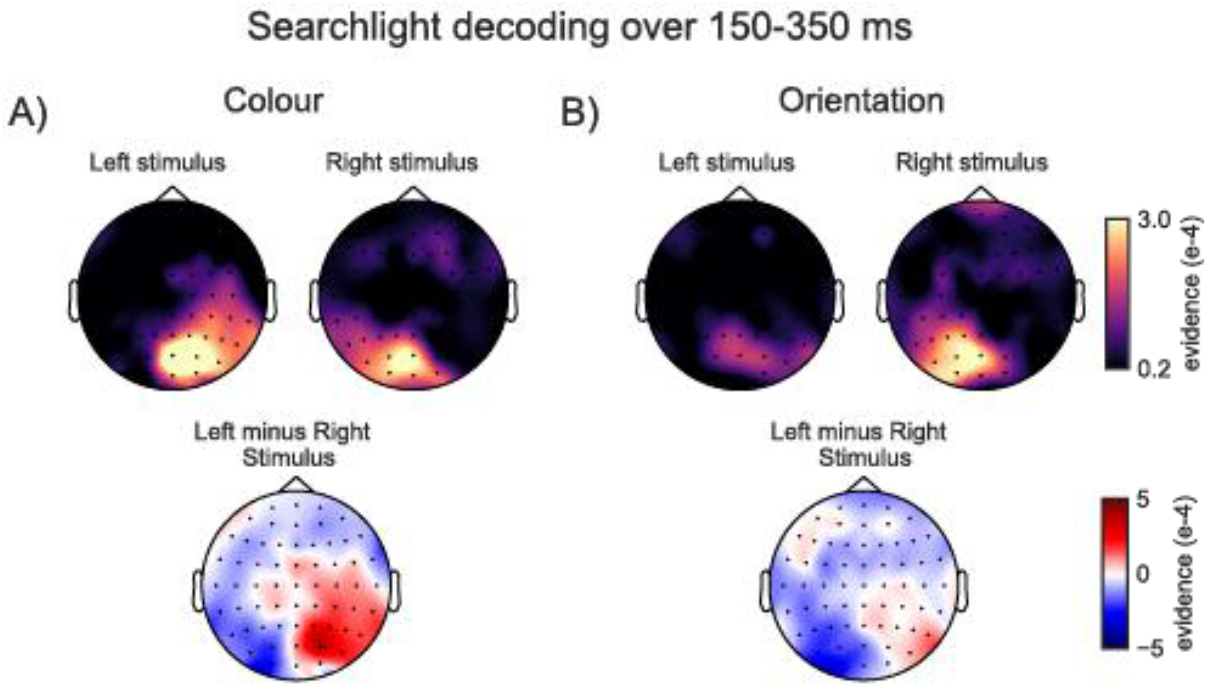
Colour and orientation decoding are primarily driven by posterior electrodes contralateral to the decoded stimulus. After averaging over data within the 150- to 350-ms time window, we ran a searchlight analysis using only the spatial pattern (in contrast to **Fig. 3** where we also incorporated spatial-temporal information). Top row topographies show which electrodes show highest mean cosine-convolved evidence for A) colours and B) orientations presented on the left and on the right. Posterior contralateral electrodes show highest evidence. Bottom panels show the difference between left and right feature decoding topographies, highlighting the lateralisation of the evidence.

Instead of concatenating data over the window of 150 ms until 350 ms post stimulus onset, we averaged data into an average pattern and proceeded with the same analyses as depicted in (**Fig. 4**). For colour bins, 11/12 colours could be decoded significantly against a *p* < .05 threshold (**Fig. S4**; mean t_29_ = 3.391; min = 0.530; max = 6.040), and for orientation all 6/12 orientations could be decoded significantly against an uncorrected single-tail threshold (mean t_29_ = 2.436; min = 0.530; max = 4.561). After applying Bonferroni correction, 5/12 colour bins were significant compared to 4/12 orientations.

**Fig. S4.**
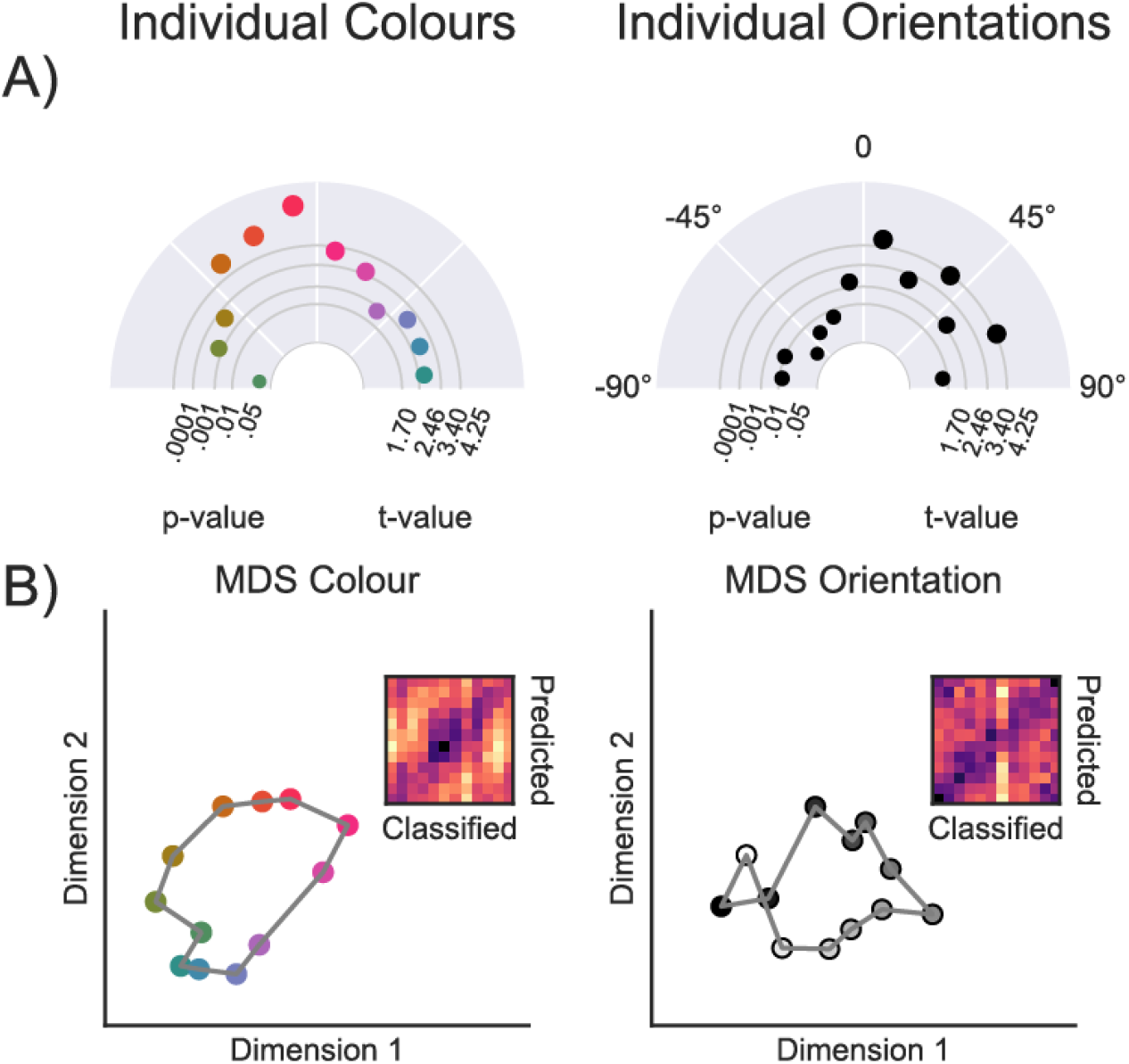
Decoding of individual features between 150 and 350 ms. A) This polar plot illustrates the decoding evidence of 12 individual colours and orientations. The distance from the centre depicts the t-value of testing the cosine-convolved tuning curve for a single feature, left and right combined, across participants, relative to zero. The size of the dots mirrors the magnitude of the evidence for that feature B) Two-dimensional visualisation of the similarity matrix in which we observed a circular configuration for colour features. Dissimilarity matrices show feature (colour or orientation) on the y-axis and distance between bins of the tuning curve on the x-axis.

**Fig. S5.**
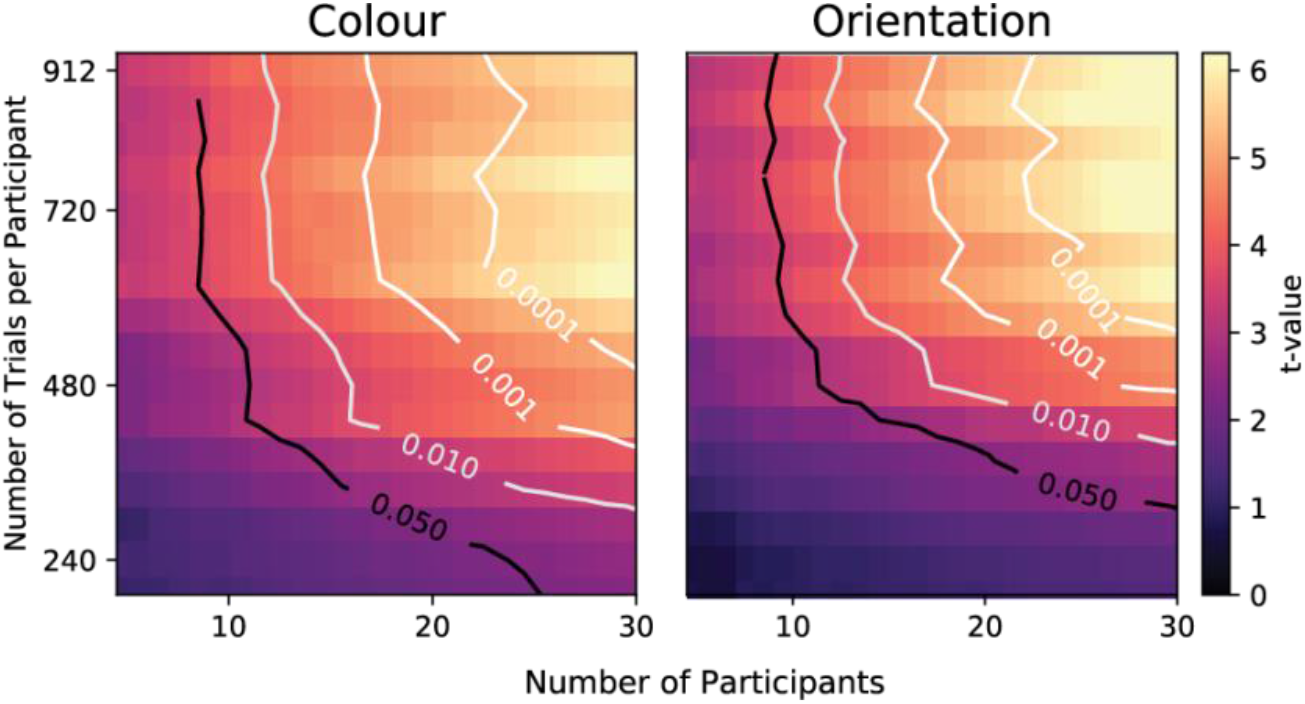
Feature decoding as a function of trial and participant numbers. Average cosine-convolved evidence between 150 and 350 ms for the average of left and right feature decoding. Contours mark the .05, .01, .001, and .0001 p-value thresholds for n-1 degrees of freedom.

